## Supplementary material for "Multiomics Analyses Reveal Dynamic Bioenergetic Pathways and Functional Remodeling of the Heart During Intermittent Fasting": Legends for Supplementary Figures, Tables, and Movies

**Supplementary Legends**

**S. Figure 1: Experimental design and details of the study cohort**

**(a)** Schema showing the experimental design of the current study. At the 6-month time point, mice were euthanized, and hearts were removed and processed for transcriptomic and proteomic analyses. Another set of mice were used for echocardiography experiments (n = 10 mice/group) **(b)** Fasting blood glucose and ketone levels at the indicated time points. Values are the mean ± SEM (n = 20 mice/group). *p < 0.05, **p < 0.01, ***p < 0.001 compared to the AL group value (one-way ANOVA with Tukey’s post hoc test). **(c)** Average monthly body weights (g) of AL and IF mice during the experiments (6 months) (one-way ANOVA with Tukey’s post hoc test). **(d)** Overall compositional calorie intake by the mice in terms of protein, fat and carbohydrates per body weight (kcal/g)

**S. Figure 2. IF-responsive proteins distribution across different fasting regimens**

**(a)** The overlap among the significantly differentially expressed proteins in each IF group compared to AL (one-way ANOVA with Dunnett’s post hoc test). **(b)** Volcano plot of terms from the MSigDB_Hallmark_2020 gene set for the enrichment of enzymes modulated with IF.  The larger and darker-colored the point, the more significantly enriched the input gene set is for the term. **(c)** Comparison of kinase modulation across different IF regimens based on significance. Red indicates increased abundance, and blue indicates decreased abundance.

**S. Figure 3: Transcriptomics-based analysis**

**(a)** Hierarchical clustering of differentially expressed mRNAs in the three IF groups compared to the AL group. Upregulated genes are indicated in red and down-regulated genes are shown in blue. The color scale represents the log10 (average FPKM + 1) value. **(b)** Principal components analysis of RNA sequencing samples from AL, IF12, IF16, and EOD groups. **(c)** Venn diagram showing the distribution of differentially expressed mRNAs from IF12, IF16, and EOD groups compared to the AL group. **(d)** Volcano plots of differentially expressed mRNAs in IF12, IF16, and EOD groups compared to the AL group. The threshold of differential expression was set at p < 0.05. Each dot represents an individual gene (blue: no significant difference; red: upregulated gene; green: down-regulated gene). Volcano plot of statistical significance (-log10 q-value) against enrichment (log2 fold change) of differentially expressed genes. **(e)** Significantly enriched GO terms in each category of the biological process for each IF group compared to the AL group. (n = 5 mice/group). FPKM, fragments per kilobase of transcript per million mapped reads.

**S. Figure 4: Comparison of IF-induced transcriptome and proteome rewiring**

**(a)** Comparison of proteome and transcriptome abundances in AL and different IF groups. Protein levels are represented by average log-transformed normalized abundances, and transcriptome levels are represented using average log-transformed FPKM within each group **(b)** Functional processes and pathways altered by changes in transcriptome and proteome abundances at IF12 (Benjamini-Hochberg FDR < 0.05, 2-D annotation enrichment using MANOVA test). Each dot on the plot represents a modulation in a pathway or process. The scores are indicative of abundance changes of transcriptome or proteome levels over AL. Upregulation and downregulation of components are denoted by positive and negative scores, respectively. **(c)** EOD induced fold changes in mRNA (x-axis) and protein abundances of enzymes (y-axis) in comparison to AL .

**S. Figure 5: Analysis of the core clock components genes**

Differentially expressed mRNAs for clock genes in the three IF groups compared to the AL group (heat map). Most of the clock gene transcripts displayed differential expression profile compared to the AL control group irrespective of the different fasting regimens (Wald test as implemented in DESeq). IF12 and IF16 groups showed similar trend and did not particularly differ from the EOD fasting group. Red border denotes increased expression, and blue border denotes decreased expression compared to AL.

**S. Figure 6: Network mapping of IF-responsive proteins across different regimens**

Proteins that showed significantly altered protein abundance in response to **(a)** IF12 (yellow), **(b)** IF16 (blue), and **(c)** EOD (red) regimen are overlaid on the consensus protein-protein interaction network constructed using all IF-responsive proteins.

**S. Figure 7: Functional clusters enriched in IF-modulated network**

Close-knit protein clusters spanning different functional processes from the IF-responsive protein network are shown. The color of each node indicates the IF regimen at which the protein was differentially expressed. Yellow, blue, and red indicate IF12, IF16, and EOD regimens, respectively.

**S. Figure 8: Association perturbations across different IF regimens.**

**(a)** Proteins with altered functional network associations (colored nodes) against a background protein association network assembled from positive and negative co-regulation analysis of all quantified proteins (p ≤ 0.05, hypergeometric test). Only those protein co-regulations that were significant (Benjamini-Hochberg adjusted p ≤ 0.01, correlation test) were included in the background protein association network. The functional involvement of perturbed proteins across different IF regimens is shown below. **(b)** Proteins showing regimen-specific association perturbations in selected biological processes.

**S. Figure 9: (a)** Increased representation of mitochondrial proteins among all association perturbed proteins (radar plot). Numbers represent significant of enrichment (negative log10 adjusted p value). **(b)** Association network of mitochondrial proteins impacted by IF in the heart. The perturbed proteins specific to each IF or AL regimen are indicated by yellow (AL), green (IF12), blue (IF16) and pink (EOD) nodes. Perturbations inferred from positive associations are indicated as red edges, and those from negative associations are shown as blue edges. All gray nodes denote proteins sharing association perturbation with the indicated mitochondrial proteins. Colored borders around the nodes are representative of pathways or processes.

**S. Figure 10: Site-specific alteration of IF-responsive phosphoproteins**

**(a)** **(b-e)** Temporal alteration of specific IF-responsive phosphorylation sites in insulin signaling (a), lipid metabolism (b), regulation of heartbeat (c), and heart structural remodeling/development (d). Red indicates increased phosphosite abundance, and blue denotes decreased phosphosite abundance. The overall trend among the diet groups based on the phosphoproteome landscape is indicated as brown triangles.

**S. Figure 11: Echocardiographic analyses**

(**a**) Reduced mitral E/A ratio (left panel) and increased deceleration time (right panel) in hearts of IF16 compared to AL mice suggest that IF enhances diastolic function. Values are the mean ± SD; n=10 each; *p < 0.05, ***p < 0.001. Representative images of mitral pulse waves are shown in the middle panel (Unpaired t-test). (**b**) Comparision of heart weight:body weight ratio between AL and IF16 groups. Values are the mean ± SEM; ****p < 0.0001; n=36-40 each.

Table S1. List of all identified proteins in heart tissues of different dietary groups.

Table S2. IF-responsive differentially expressed proteins in heart tissues among different dietary groups.

Table S3. Functional processes and pathways enriched among different IF groups.

Table S4. RNA-seq based transcriptome profiling for heart tissues obtained from mice in different IF regimens.

Table S5. Functional processes and pathways differentially enriched in proteome and transcriptome.

Table S6. Node properties of time-resolved IF protein network.

Table S7. Perturbation of protein association co-regulation in different dietary groups.

Table S8. Quantitative phosphoproteome of heart tissues across different dietary groups.

Table S9. Echocardiographic assessment of myocardial functional and cardiac dynamics.

Movie S1.

Baseline *ad libitum* echocardiographic measurements.

Movie S2.

Baseline *intermittent fasting* echocardiographic measurements.

Movie S3.

Dobutamine stress *ad libitum* echocardiographic measurements.

Movie S4.

Dobutamine stress *intermittent fasting* echocardiographic measurements.
