## Supplementary figures and images for "Multiomics Analyses Reveal Dynamic Bioenergetic Pathways and Functional Remodeling of the Heart During Intermittent Fasting"

### Supplementary Figure 1

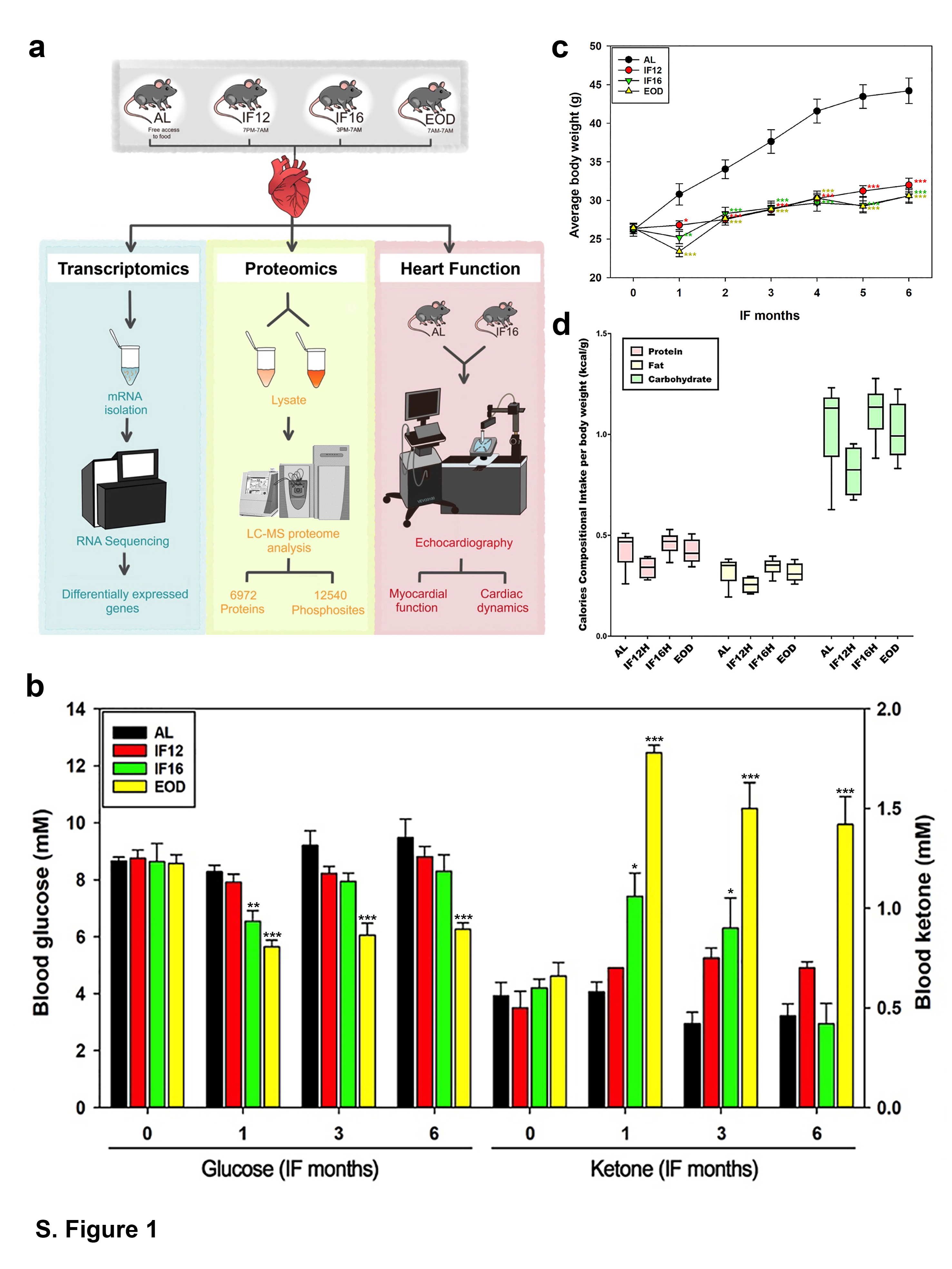

### Supplementary Figure 2

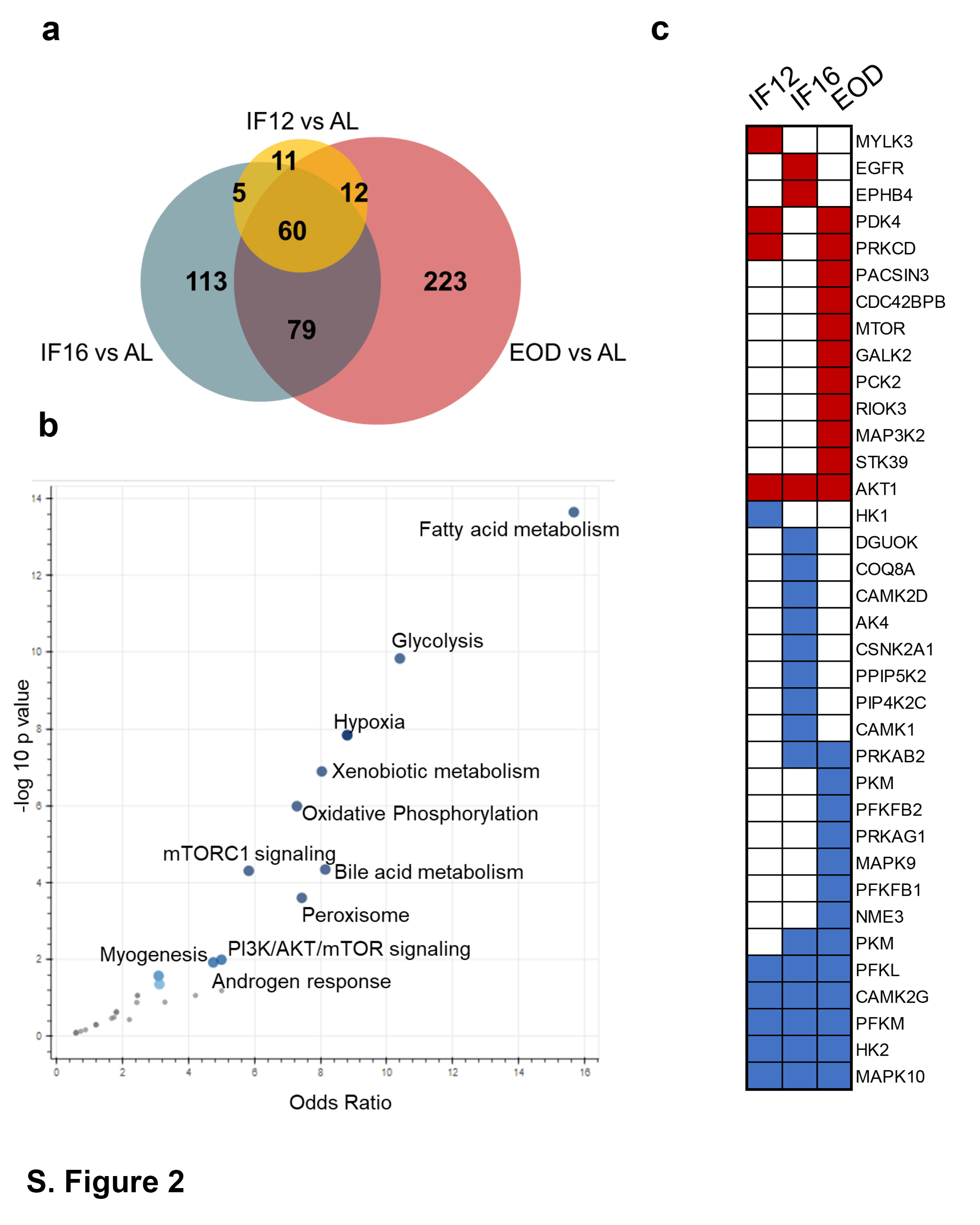

### Supplementary Figure 3

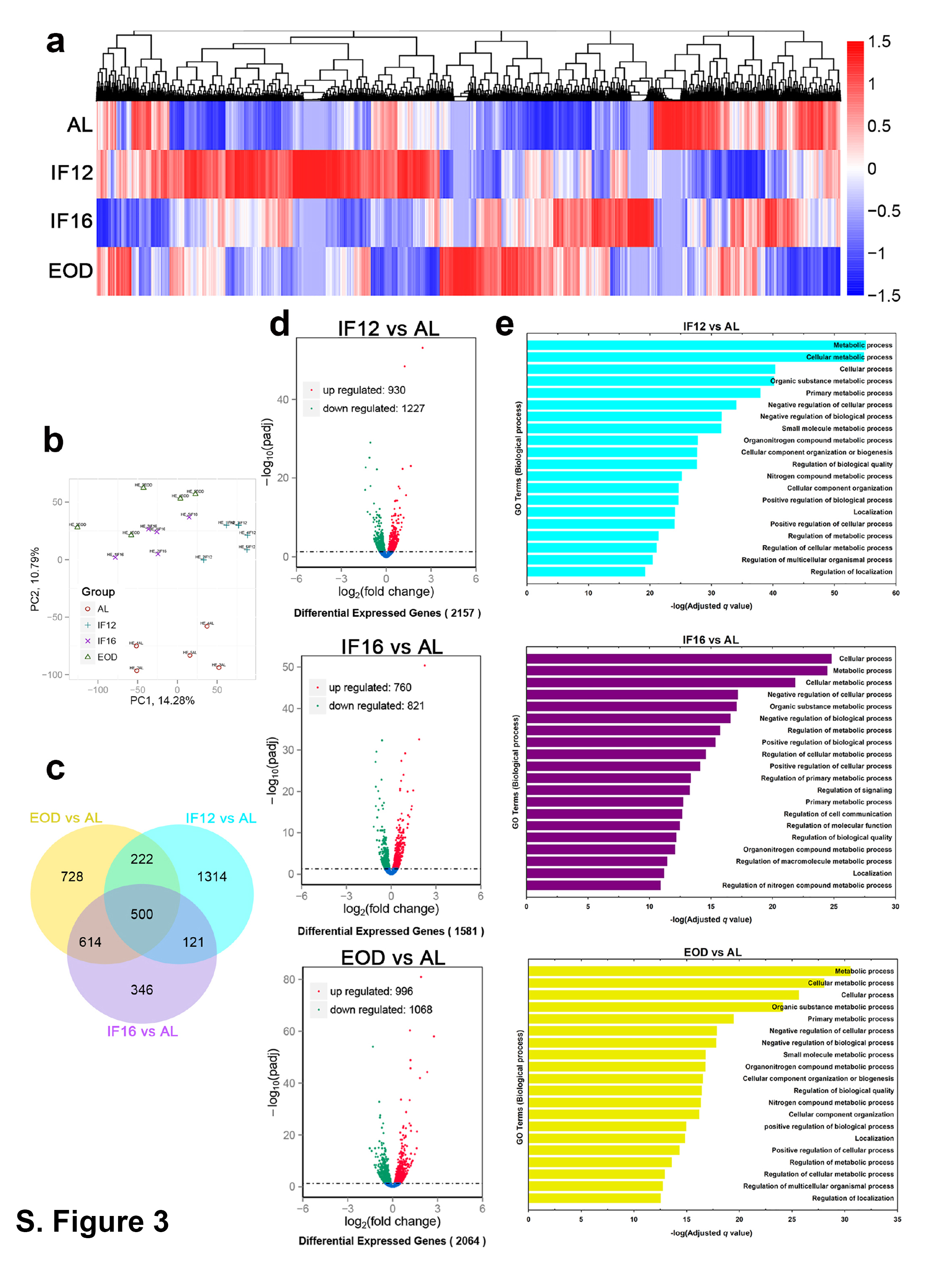

### Supplementary Figure 4

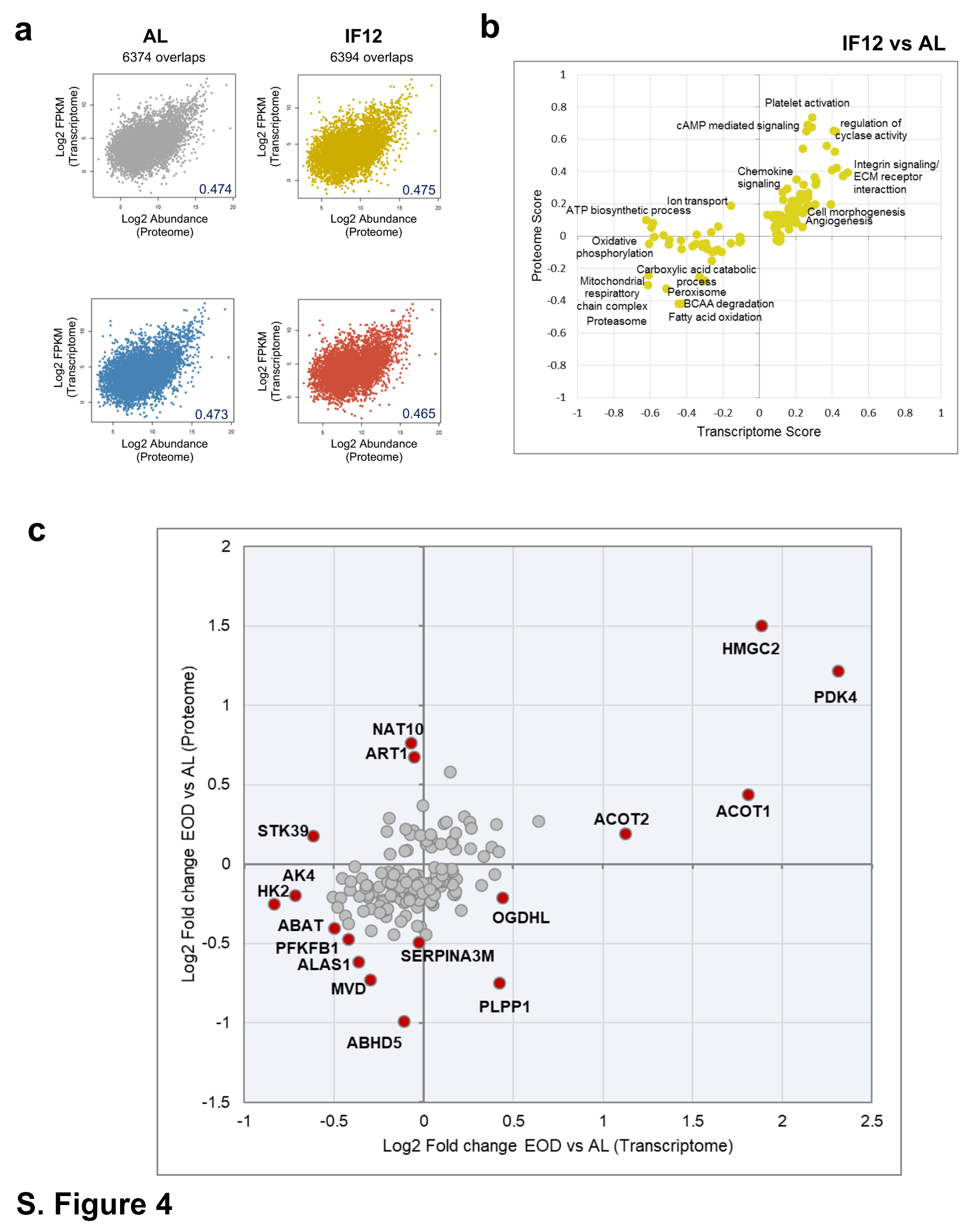

### Supplementary Figure 5

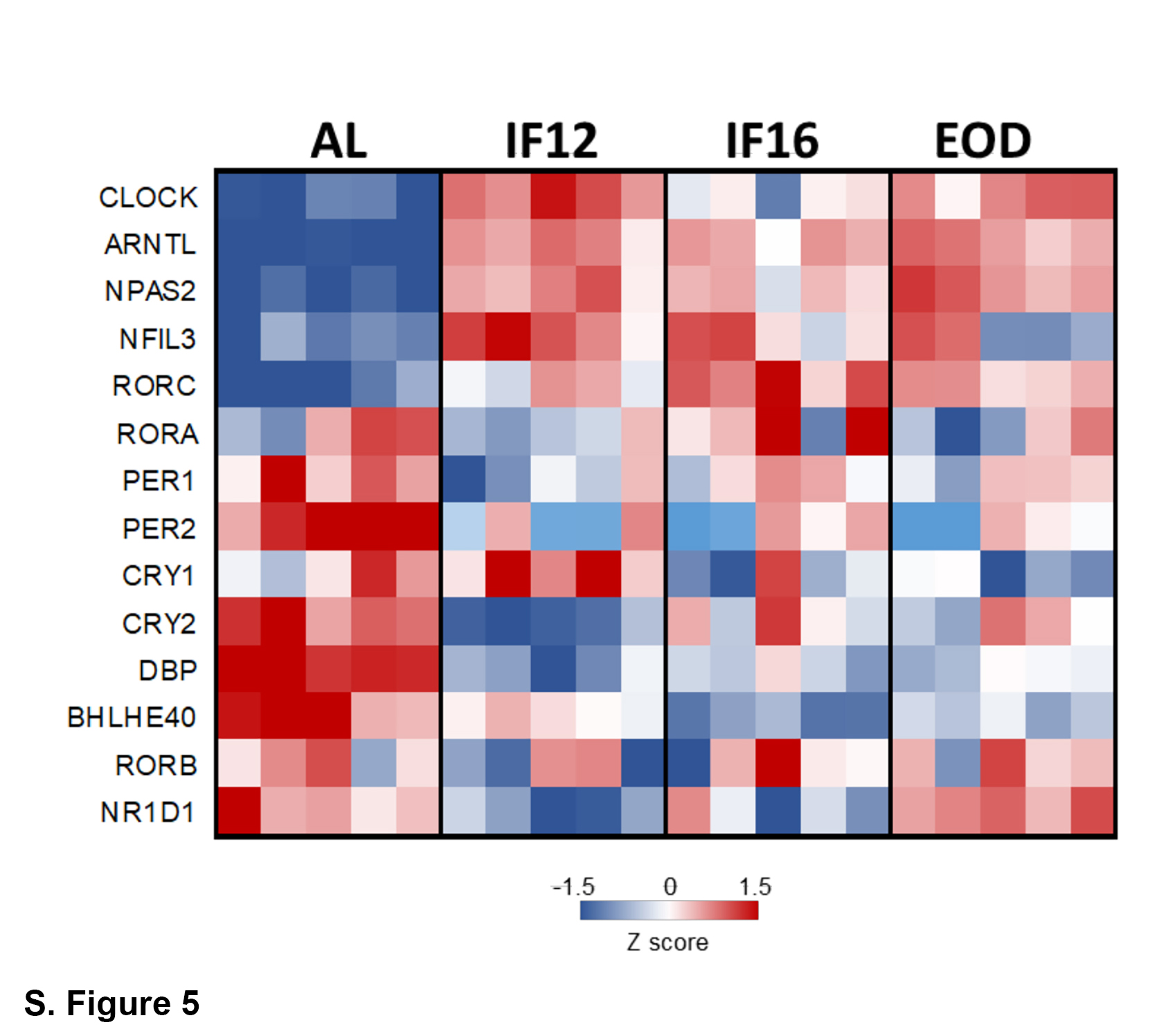

### Supplementary Figure 6

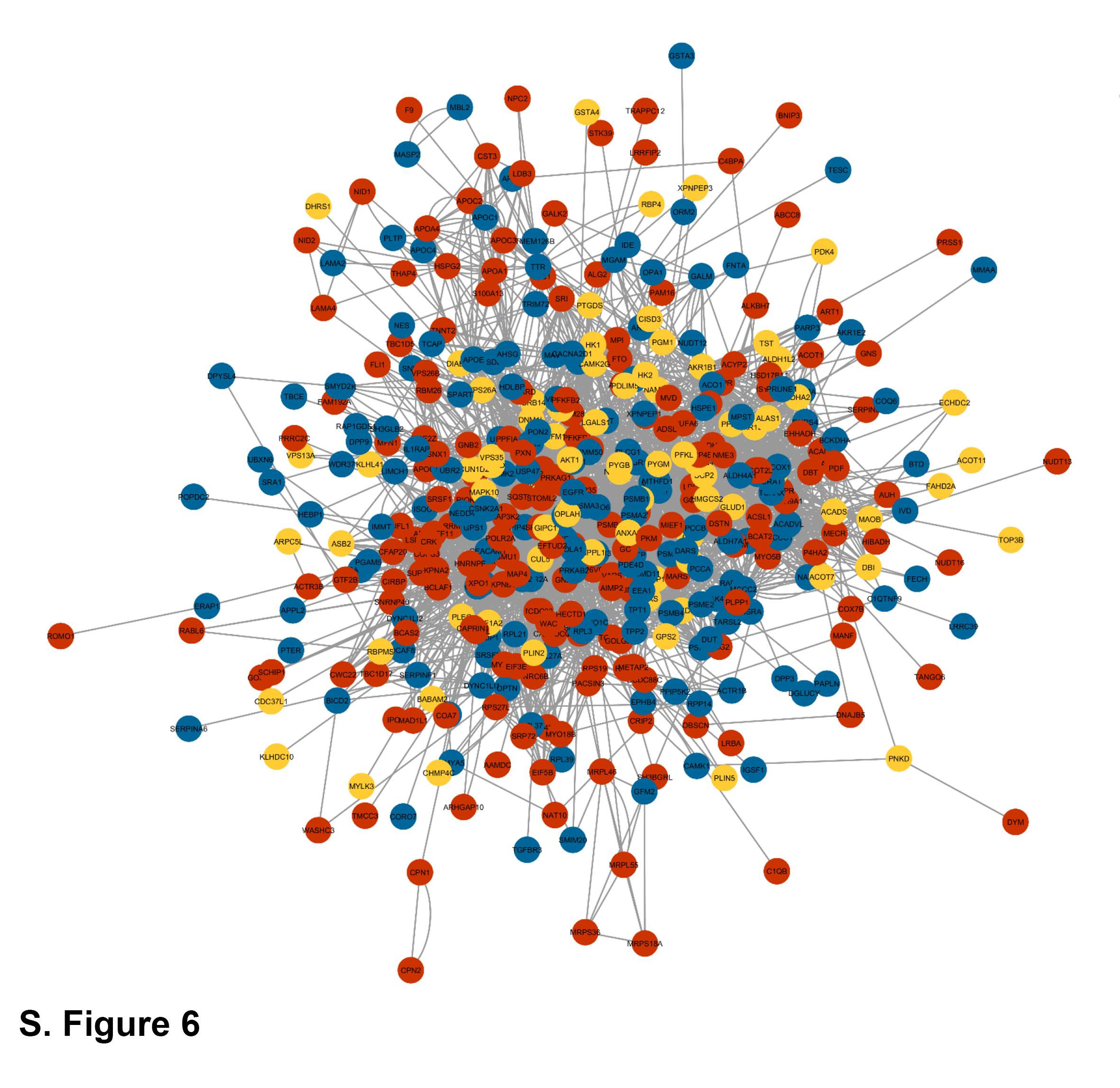

### Supplementary Figure 7

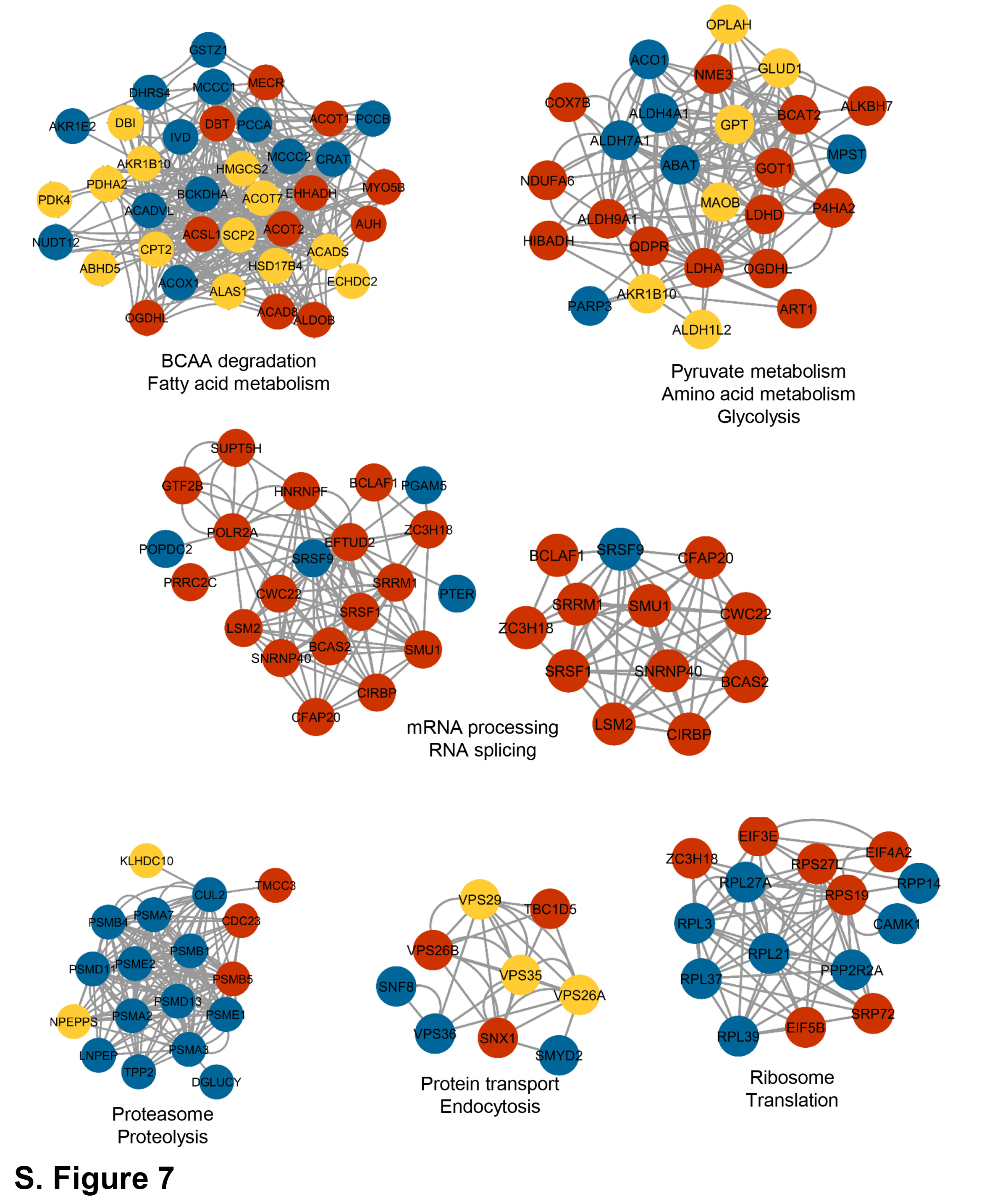

### Supplementary Figure 8

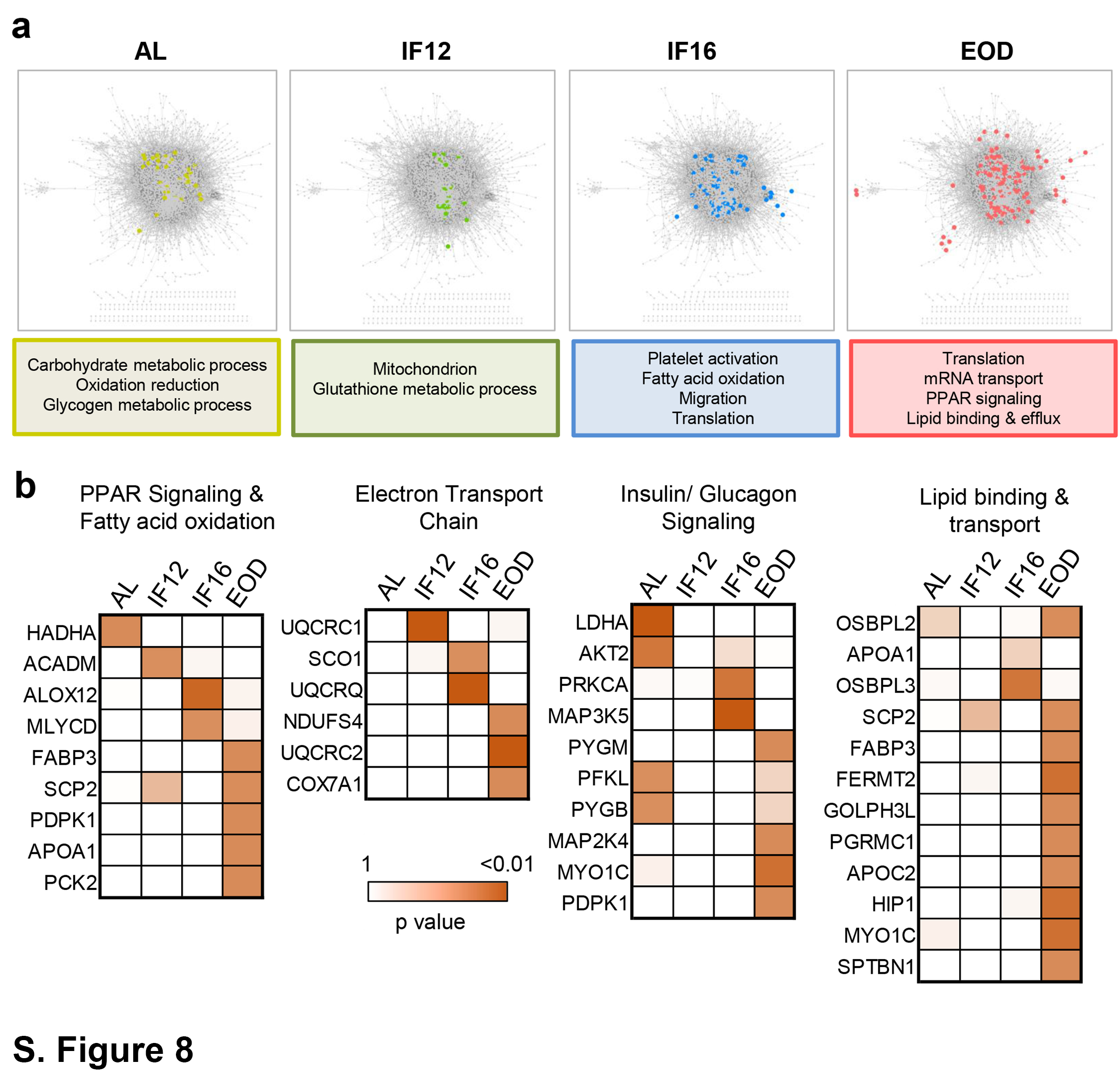

### Supplementary Figure 9

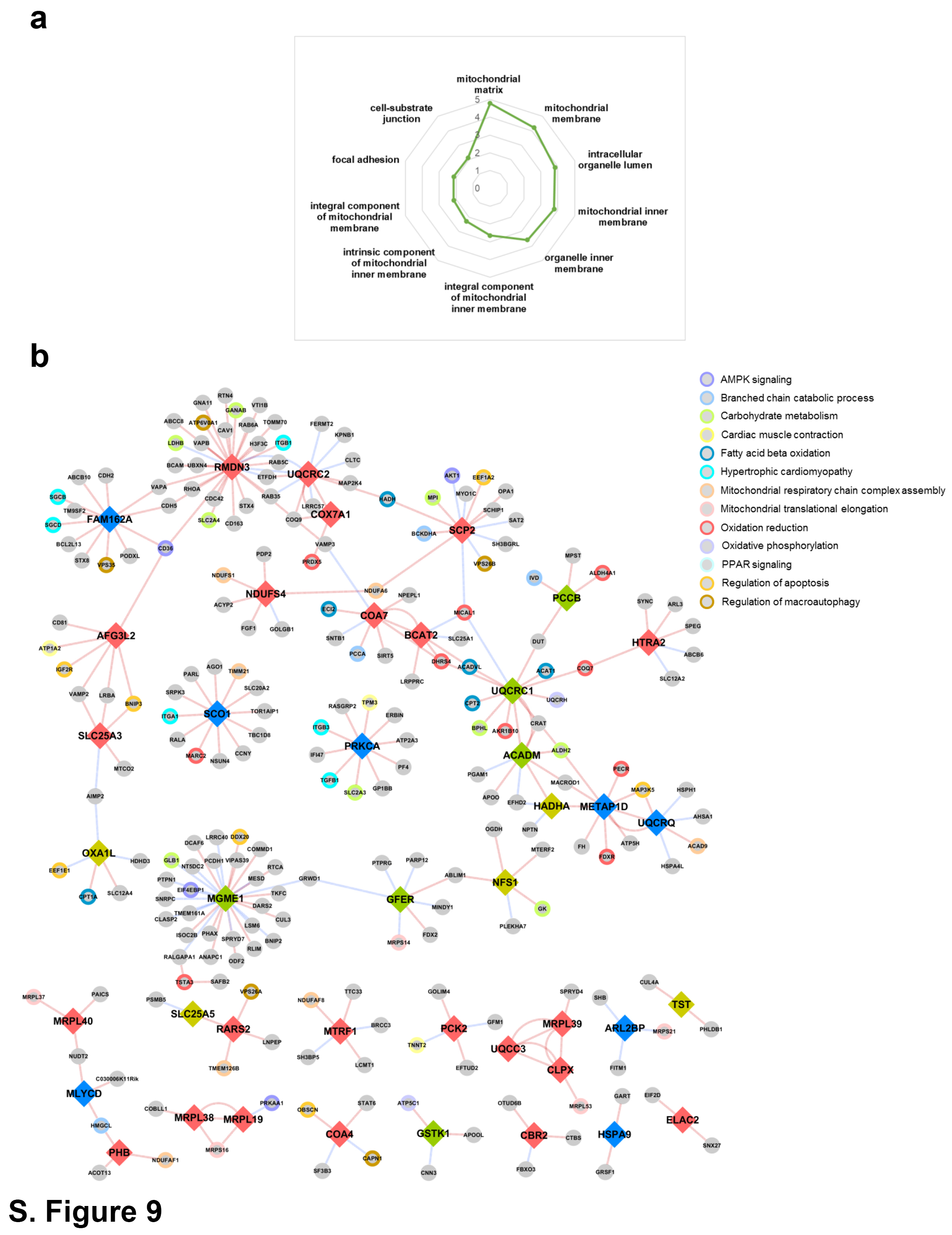

### Supplementary Figure 10

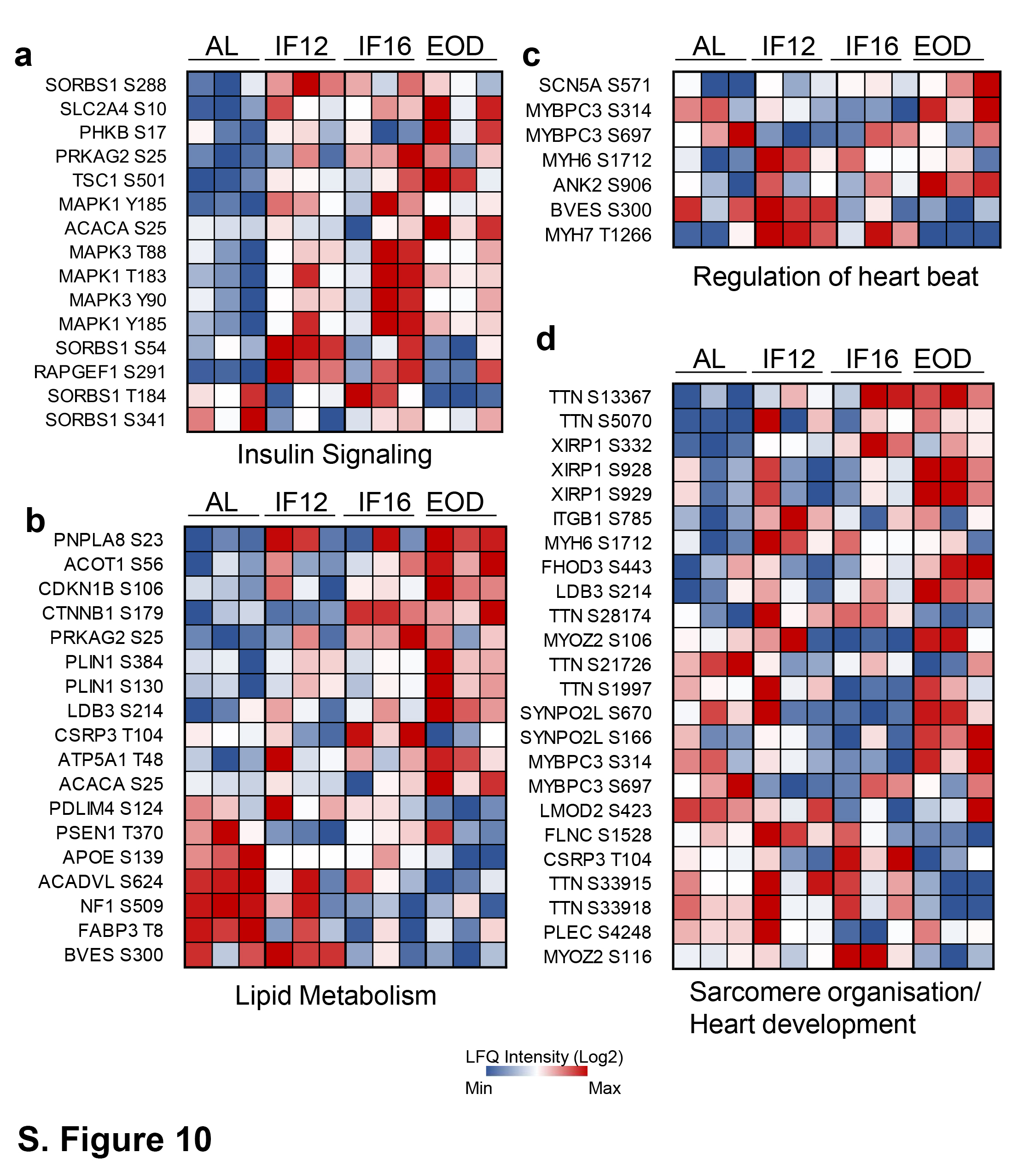

### Supplementary Figure 11

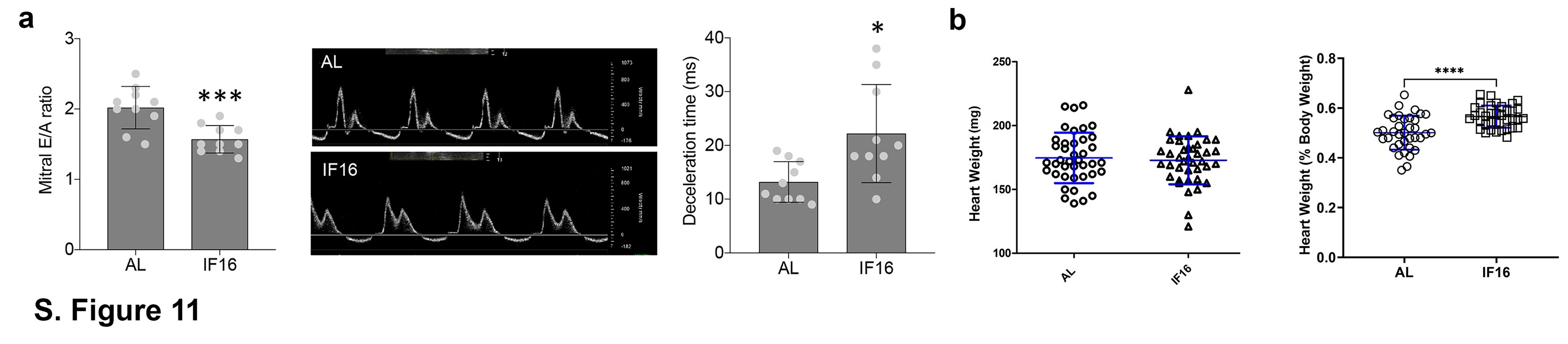
